# A Multifactorial Model of T Cell Expansion and Durable Clinical Benefit in Response to a PD-L1 Inhibitor

**DOI:** 10.1101/231316

**Authors:** Mark DM Leiserson, Vasilis Syrgkanis, Amy Gilson, Miroslav Dudik, Sharon Gillett, Jennifer Chayes, Christian Borgs, Dean F Bajorin, Jonathan Rosenberg, Samuel Funt, Alexandra Snyder, Lester Mackey

## Abstract

Checkpoint inhibitor immunotherapies have had major success in treating patients with late-stage cancers, yet the minority of patients benefit [1]. Mutation load and PD-L1 staining are leading biomarkers associated with response, but each is an imperfect predictor. A key challenge to predicting response is modeling the interaction between the tumor and immune system. We begin to address this challenge with a multifactorial model for response to anti-PD-L1 therapy. We train a model to predict immune response in patients after treatment based on 36 clinical, tumor, and circulating features collected prior to treatment. We analyze data from 21 bladder cancer patients [2] using the elastic net high-dimensional regression procedure [3] and, as training set error is a biased and overly optimistic measure of prediction error, we use leave-one-out cross-validation to obtain unbiased estimates of accuracy on held-out patients. In held-out patients, the model explains 79% of the variance in T cell clonal expansion. This predicted immune response is multifactorial, as the variance explained is at most 23% if clinical, tumor, or circulating features are excluded. Moreover, if patients are triaged according to predicted expansion, only 38% of non-durable clinical benefit (DCB) patients need be treated to ensure that 100% of DCB patients are treated. In contrast, using mutation load or PD-L1 staining alone, one must treat at least 77% of non-DCB patients to ensure that all DCB patients receive treatment. Thus, integrative models of immune response may improve our ability to anticipate clinical benefit of immunotherapy.

## Introduction

Immunotherapies such as checkpoint inhibitors have become a major success in treating patients with late-stage cancers, in many cases leading to durable responses [1,4–7]. The basis for this success is thought largely to result from the somatic mutations present in cancer cells allowing the immune system’s T cells to distinguish cancer from normal cells, in part because mutations may lead to the presentation of neoantigens on the cancer cell surface by the major histocompatibility complex [8,9]. However, many cancers develop mechanisms for suppressing the immune system, including expression of checkpoint molecules [1]. The promise of checkpoint inhibitor cancer therapies is predicated on counteracting checkpoint molecules to unleash the immune system to selectively kill cancer cells.

Despite checkpoint inhibitors’ unprecedented successes, there is an urgent need to improve prediction of patient response to checkpoint inhibitor immunotherapy. Response rates vary across patients, and known biomarkers for response such as high mutation load are not predictive for every patient [7,10–13]. Several studies have refined definitions of mutation load and assessment of neoantigen quality to improve prediction of response [8,14], but this process remains imperfect. Thus, predicting response is critical for identifying patients who are likely or unlikely to benefit, anticipating adverse responses to treatment [15], and accelerating the development of new treatments. Further, effective models for predicting response may point to molecular features that can be measured and monitored through non-invasive methods.

A key challenge for predicting response is modeling features of the immune system and cancer simultaneously. Recently, clinicians have begun to collect a wealth of molecular tumor and immune system data before and during immunotherapy, but researchers have yet to model how molecular and clinical features interact to affect response.

To address this challenge, we develop a multifactorial model for response to checkpoint inhibitors. Our approach uses the elastic net [3] -- a machine learning method for regression that automatically selects informative features from the data -- and models clinical, tumor, and immune system features simultaneously. We applied our model to the data of Snyder et al. [2], who measured mutations and gene expression in the tumor and T cell receptor (TCR) sequences in the tumor and peripheral blood in urothelial cancers treated with anti-PD-L1. Rather than model the clinical response of each patient directly, we modeled the response of each patient’s immune system and used the predicted immune responses to stratify patients based on expected clinical benefit. By modeling the immune response, we have the advantage of predicting fine-grained, molecular measurements that are associated with clinical response.

## Methods

All of our analyses were conducted in Python 3 using open source software, and Jupyter notebooks that replicate our experiments are publicly available at https://github.com/lrgr/multifactorial-immune-response.

### Patient Data

We used the patient data collected by Snyder et al. [2]. For the data collection details and the Institutional Review Board approval see [2].

### Multifactorial modeling of clonal expansion

In our first analysis, we develop a predictive model of the log number of tumor-infiltrating lymphocyte (TIL) clones that expanded in the blood three weeks after each patient’s initial immunotherapy treatment. We chose to model TIL clone expansion, as it is a finer-grained and more immediate measurement of patient response than standard clinical data that still exhibits positive association with durable clinical benefit [2]. Our analysis is based on the 21 patients with recorded clonal expansion, whole exome sequencing (WES), and RNA sequencing (RNA-seq) data.

Our predictions were based upon 36 patient features derived from nineteen attributes of each patient collected prior to treatment (see Table 1). For the one patient who did not have a 5-factor score [16] recorded, because atezolizumab was given as first line therapy, we substituted the patient’s Bajorin risk score [16]. We encoded Prior BCG and Albumin < 4 as binary features, with values in {0,1}. For each of the remaining attributes, we included both the raw feature value, x, and a log(1+x) transform of the feature value as inputs to our models to capture nonlinear relationships between inputs and the target.

**Table 1:** Patient attributes collected prior to treatment and processed as learning pipeline inputs.

| Clinical | Tumor | Circulating |
| --- | --- | --- |
| Prior intravesical Bacillus Calmette–Guérin (BCG) | Missense SNV count (mutation load) | Productive unique TCR count |
| Age | Expressed missense SNV count | Clonality (TCR) |
| Albumin < 4 | Neoantigen count | Diversity (TCR) |
| Baseline neutrophil to lymphocyte ratio | Expressed neoantigen count | T cell fraction |
| Time since last chemotherapy (days) | Clonality (TCR) | Top clone frequency (%) |
| 5-factor score [16] | Diversity (TCR) |  |
| Number of chemotherapy regimens | T cell fraction |  |

We trained two machine learning models to predict log clonal expansion from our input features: the elastic net [3], a high-dimensional linear regression procedure designed to reduce overfitting and be robust to irrelevant features, and random forests [17], a highly-nonlinear regression procedure based on averaging the predictions of many randomly constructed decision trees. We fit the elastic net using the Python scikit-learn [18] ElasticNetCV function with feature normalization, candidate L_1_ ratio hyperparameters [.1, .5, .7, .9, .95, .99], hyperparameters selected using leave-one-out cross-validation, tolerance 10^−7^, and a maximum number of iterations of 10^6^. We fit the random forest model using the Python scikit-learn RandomForestRegressor with 1000 decision trees. Prior to fitting each regression model, missing input feature values were imputed using the median non-missing feature value in the training set.

### Leave-one-out error analysis of expansion predictions

To estimate the effectiveness of our learning pipelines at predicting the clonal expansion of new patients, we conducted a leave-one-out cross-validation (LOOCV) analysis. Specifically, for each of the 21 patients in turn, we withheld that individual’s data from the training set, fit each of our learning pipelines (including hyperparameter selection) on the remaining 20 patients, and formed a prediction of the held-out patient’s log clonal expansion using each of the learned models. We then compared each learned model’s predictions with the observed log clonal expansion for the held-out patient and computed the squared error. By computing the average of this held-out squared error across all patients, we obtain an unbiased estimate of each learning pipeline’s error in predicting previously unseen patient expansion. Importantly, we do not use training set error as a measure of predictive performance in any of our analyses, as such in-sample error is known to provide biased, overly optimistic estimates of predictive performance. Saria et al. [19] use a similar leave-one-out analysis to assess predictive models of preterm infant illness. We moreover compute a measure of variance explained in held-out patients by computing one minus the ratio of the LOOCV mean squared error to the empirical variance of log clonal expansion. Finally, following [20], we conduct a nonparametric test of association between the input features and clonal expansion based on the pipeline’s LOOCV error. Specifically, we perform a permutation test that compares the observed LOOCV error to the distribution of LOOCV errors obtained when patient immune responses are permuted in our cohort uniformly at random. The null hypothesis of no association between input features and clonal expansion is rejected whenever the observed LOOCV error is unusually small (i.e., when the leave-one-out prediction accuracy of the learning pipeline is unusually high).

### Feature importance

To assess the degree to which different classes of features contribute to the predictive accuracy of our learning pipelines, we assigned each input feature to a category (‘Clinical’, ‘Circulating’, or ‘Tumor’) and, for each category, repeated our LOOCV analysis with features from that category excluded from the model.

In addition, inspired by [17,20], we test for association between clonal expansion and the input features belonging to a given category when the remaining input features are also available to the model. To achieve this, we perform a permutation test comparing observed LOOCV error when all features are presented to the model to the distribution of LOOCV errors obtained when the vector of feature values belonging to a given category are permuted in our cohort uniformly at random. The null hypothesis of no association between clonal expansion and the input features belonging to a category is rejected whenever the observed LOOCV error is unusually small.

### Implications for durable clinical benefit

Snyder et al. [2] previously demonstrated a positive association between the number of TIL clones that expanded in the blood two weeks after treatment and the durable clinical benefit of cancer immunotherapy. Here, a treatment is said to have durable clinical benefit (DCB) for a patient if the patient experiences progression free survival for at least six months after treatment. To assess whether our learning pipelines are also predictive of durable clinical benefit for previously unseen patients, we compare the distribution of held-out expansion predictions for those patients who did and did not experience DCB.

## Results

### Leave-one-out error analysis of expansion predictions

We used the elastic net [3], a machine learning method that automatically selects informative features from the data, to model the relationship between immune response and the clinical, tumor, and circulating features from Snyder et al. [2]. On our dataset, a baseline prediction rule that ignores all patient features and predicts the mean log clonal expansion for all patients achieves a mean squared error (MSE) of 0.838. This baseline level of error represents the total variance of log clonal expansion that may be explained by patient features. By leveraging patient features, the elastic net model achieves a large reduction in prediction error, explaining 79% of this variance in held-out patients with a leave-one-out cross-validation (LOOCV) MSE of 0.176. We reiterate that we do not use training set error to evaluate our learning pipelines, as such in-sample error is known to provide biased, overly optimistic estimates of prediction error.

Rather, our assessments are based on patients withheld from the training set using leave-one-out cross-validation (LOOCV) to provide unbiased estimates of predictive performance. Figure 1a highlights the accuracy of the elastic net model by plotting elastic net predictions against the ground truth clonal expansions of each held-out patient; perfect predictions would lie on the red line. Permutation testing with 1000 random permutations of patient responses demonstrated a statistically significant association between patient features and clonal expansion evidenced by the small elastic net LOOCV error (*p* < 0.002, Figure 1b). The random forest model performed far worse with a LOOCV MSE (0.886) that exceeded the baseline MSE, indicating severe overfitting. For this reason, we focused on the elastic net learning procedure in the remainder of our analyses.

**Figure 1:**
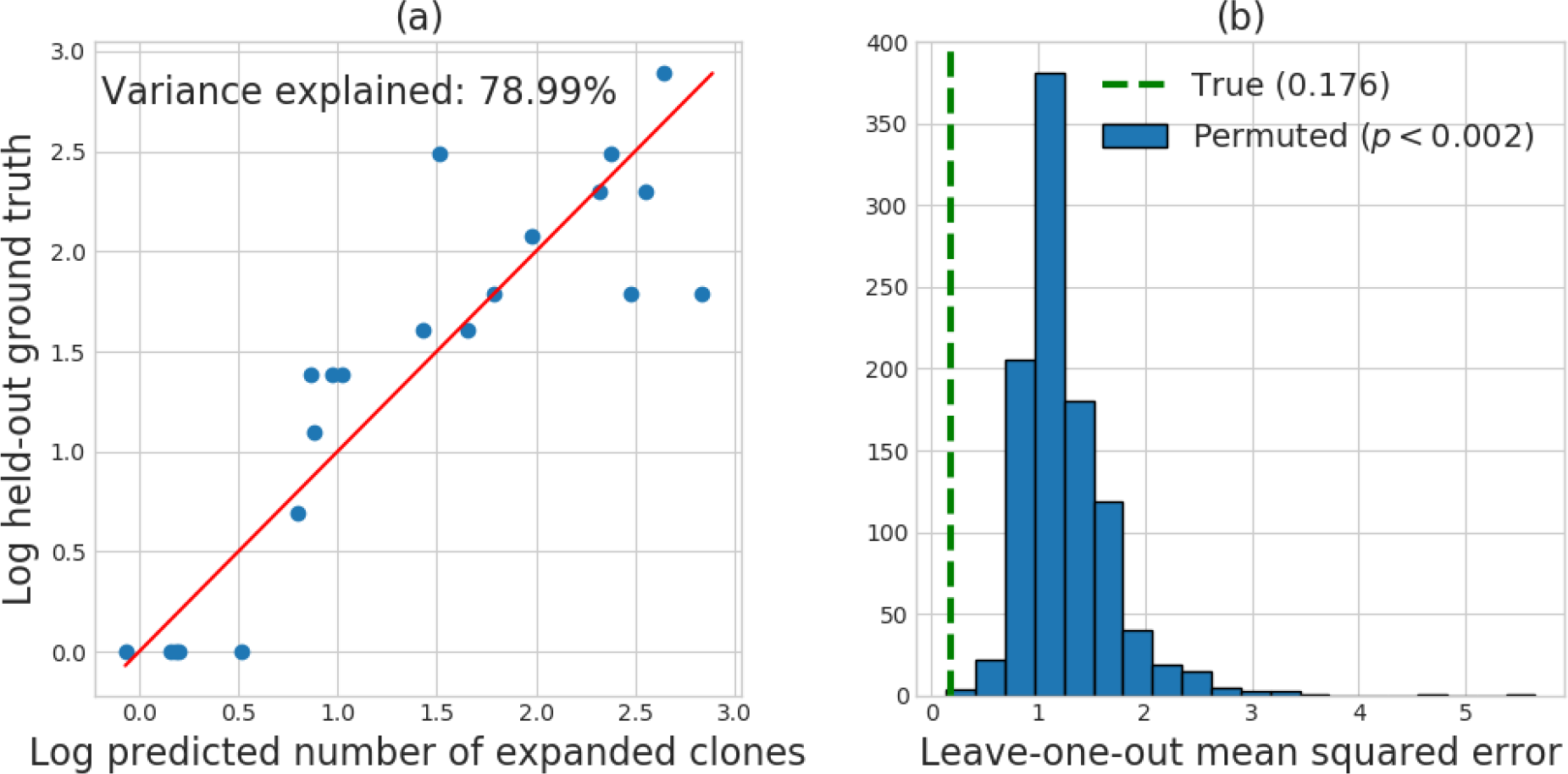
(a) Predicted log TIL expansion versus ground-truth log TIL expansion for patients held out using LOOCV. Predictions are formed using the elastic net. (b) Histogram of LOOCV error when patient responses are permuted uniformly at random 1000 times. The overlaid dotted line displays the LOOCV error obtained on the original dataset.

### Feature importance

The elastic net procedure automatically performs feature selection by setting the coefficients of some input variables to zero. The final elastic net model fit to the entire training set retains 20 of the 36 input features, a mix of clinical, tumor, and circulating patient attributes (see Figure 2). When we repeat our LOOCV error analysis using only clinical and circulating features (that is, excluding all tumor features), the variance explained in held-out patients drops from 79% to 23%. When we repeat the LOOCV error analysis using only clinical and tumor features (excluding all circulating features), the explained variance drops to 8.5%. Finally, when we repeat the LOOCV error analysis using only circulating and tumor features (excluding all clinical features), the learned prediction rules are accurate on training data but do not generalize to held-out patients. Due to this "overfitting" phenomenon, the learning pipeline without clinical features performs worse than the baseline prediction of the mean, achieving a nominal variance explained of 0%. Permutation testing with 1000 random permutations of patient features belonging to a given category demonstrated a statistically significant conditional association between clinical features and clonal expansion (*p* < 0.002), between circulating features and clonal expansion (*p* < 0.001), and between tumor features and clonal expansion (*p* < 0.004), evidenced by the small LOOCV error of the elastic net. These findings indicate that clinical, tumor, and circulating features are all contributing significantly to the predictive accuracy of the elastic net learning pipeline.

**Figure 2:**
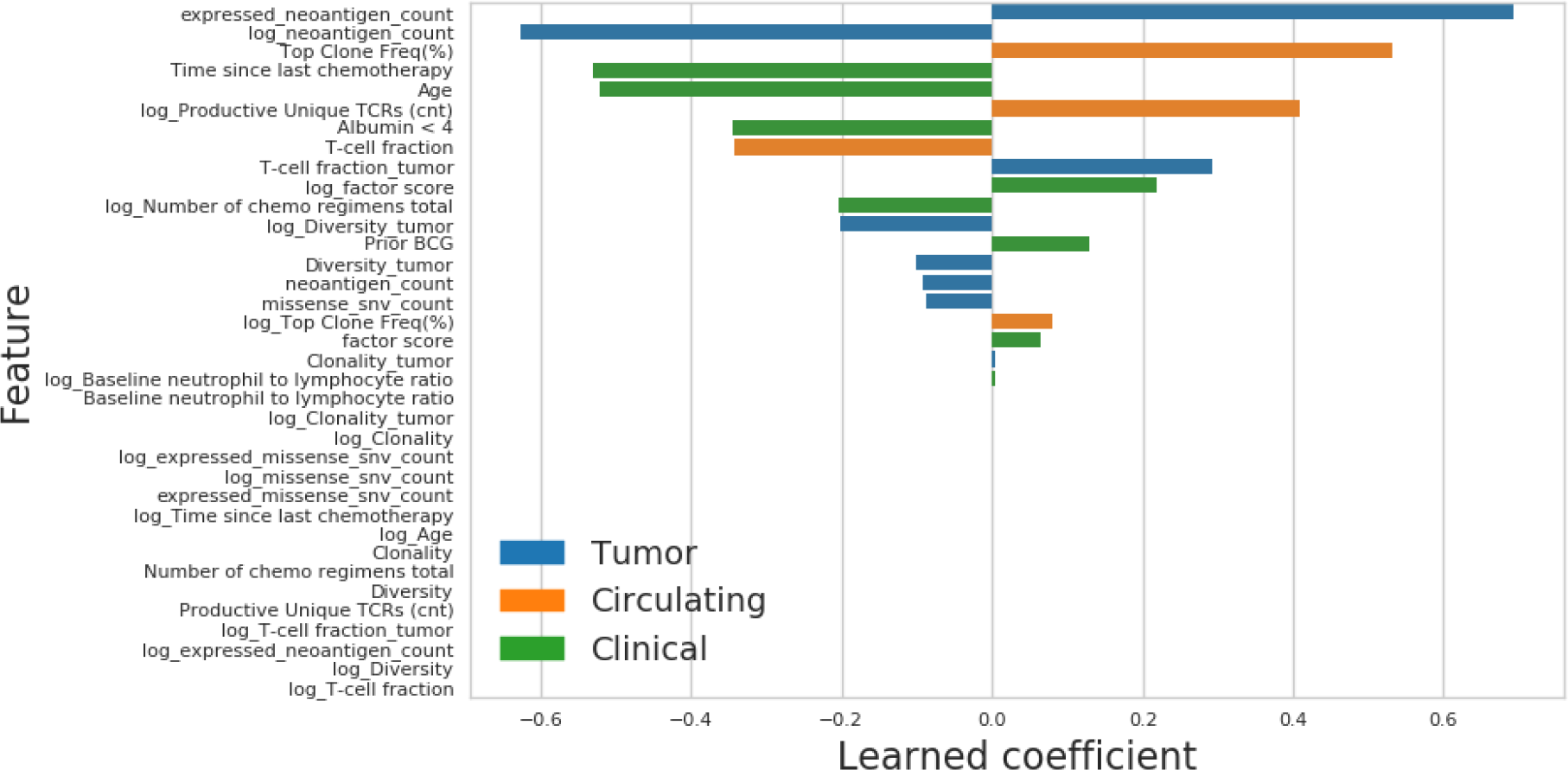
Learned elastic net coefficients and feature types.

### Implications for durable clinical benefit

While mutation load (missense SNV count) and PD-L1 staining have moved forward as clinical biomarkers, expansion of tumor-associated T-cell clones in the peripheral blood, hereafter referred to as TIL clone expansion, provides additional insight into clinical response. Figure 3a compares the distributions of the predicted number of expanded TIL clones for held-out patients who did and did not experience durable clinical benefit (DCB). This display should be contrasted with the DCB-stratified distributions of standard biomarkers like missense single-nucleotide variant (SNV) count, expressed neoantigen count, and PD-L1 staining (Figures 3b-d, respectively). Notably, 100% of patients who experienced durable clinical benefit have predicted expansion scores above the sixty-second percentile prediction for patients without durable clinical benefit. This indicates that, if patients are triaged according to predicted expansion, only 38% of non-DCB patients need be treated to ensure that 100% of DCB patients are treated. In contrast, using PD-L1 staining alone, one must treat at least 77% of non-DCB patients to ensure that all DCB patients receive treatment. Using missense SNV count or expressed neoantigen count alone, one must treat at least 85% of non-DCB patients to ensure that all DCB patients are treated.

**Figure 3:**
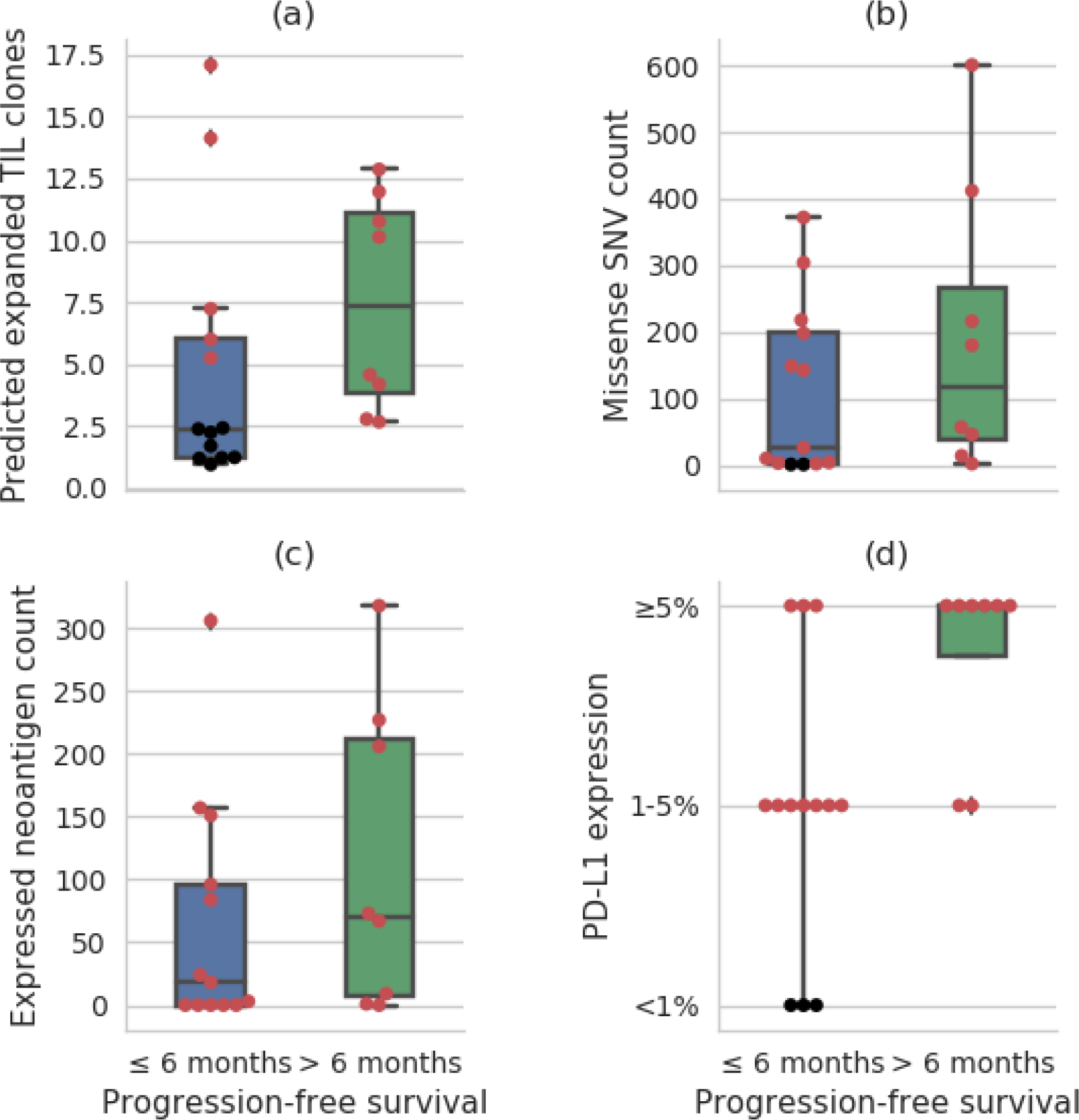
Distributions of biomarker values in patients with and without durable clinical benefit (DCB, defined as ≥ 6 months of progression-free survival): (a) predicted number of expanded TIL clones; (b) missense SNV count; (c) expressed neoantigen count; and, (d) percentage of tumor infiltrating immune cells found to be PD-L1-positive. When each biomarker alone is used for triage, the patients highlighted in red must be treated to ensure all DCB patients are treated.

Each of these biomarkers is known to be associated with DCB in cancer immunotherapy patients [21], and, indeed, mutation load and PD-L1 expression are both correlated with radiographic response in the larger urothelial cancer dataset including these patients [22]. However, none of these biomarkers alone is a perfect discriminator of DCB and non-DCB patients. The elastic net learning pipeline is able to integrate mutation load, expressed neoantigen count, and multiple other patient features into a composite biomarker with greater DCB discriminating ability.

## Discussion

We have introduced a multifactorial model for predicting response to checkpoint inhibitor immunotherapy. Our model integrates tumor, clinical, and immune features to predict a measure of immune response. Notably, we train our model to predict a fine-grained intermediate measure of immune response, the expansion of tumor-associated TCRs in the peripheral blood, which is associated with the coarser-grained clinical response, here measured as stable disease or better for at least 6 months.

We demonstrate and evaluate our model on a dataset of urothelial cancers from Snyder et al. [2]. We find that, in held-out patients, our model can predict the number of tumor infiltrating T cell clones that expand in the blood post-therapy with high estimated accuracy. In addition, if the patients in our cohort are triaged according to held-out predicted expansion, only 38% of non-DCB patients need be treated to ensure that 100% of DCB patients are treated. Moreover, we find that our model achieves the highest LOOCV accuracy when tumor, circulating, and clinical features are all included, demonstrating that they provide complementary information. We next intend to validate our model fully out-of-sample on a new cohort of urothelial cancer patients.

We anticipate that integrative models of tumor, immune system, and clinical features such as those introduced here will be necessary to understand the complexity of the anti-tumor immune response. Indeed, other groups have integrated radiographic tumor burden and TIL clone expansion to assess response to anti-PD-1 therapy in melanoma and found that responses depend on both immune and tumor features [23]. Non-invasive measurement of immune system activity and cancer genomics in the blood has the potential to transform many areas of cancer care, including early detection [24] and monitoring response [25]. When taken together, these data offer a tantalizing early look at how predictive models of the peripheral immune system, and predictive models of clinical response from peripheral immune system features, will play a role in helping to personalize immunotherapy treatment strategies and drug discovery.

## Acknowledgments

We sincerely thank Arnold Levine and Stand Up To Cancer for facilitating this collaboration.

